# Epithelial cell fusion is required for tissue repair following UV-A irradiation

**DOI:** 10.1101/2025.09.26.678815

**Authors:** Minqi Shen, Lillie G. Mitchell, Lydia W. Boer, Vicki P. Losick

## Abstract

Cell cycle-dependent and independent mechanisms lead to the generation of mononucleated and multinucleated, polyploid cells. The more than doubling of a cell’s nuclear genome by endoreplication has been found to be an adaptation to genotoxic stress, enabling cell survival despite DNA damage. However, it remains unknown whether cells that increase ploidy via multinucleation also arise in response to genotoxic stress. Here, we use ultraviolet light A (UV-A) to induce permanent DNA damage in cells within the adult fruit fly epithelium. UV-A irradiation causes an injury-like response where giant multinucleated, polyploid cells arise following cell death. The epithelial cells undergo endoreplication, which is required to restore tissue mass, but is surprisingly dispensable for tissue repair. UV-A irradiation also induces cell fusion, which generates multinucleated cells that encompass almost the entire epithelial area post injury. Cell fusion can be inhibited by expression of a dominant negative Rac or Cdc42 GTPase, which then blocks epithelial tissue repair post irradiation. Apoptotic nuclei were detected at the site of cell junction breakdown suggesting that apoptosis itself or an apoptotic signal is required for polyploidization in this model. Expression of the effector caspase inhibitor, p35, led to inhibition of apoptosis, the endocycle, and cell fusion post UV-A. Therefore, we have discovered that caspase activation is necessary for polyploidization post injury and enhancing cell ploidy via multinucleation is another strategy to enable cell survival and tissue repair following genotoxic stress.

**Significance Statement:** Polyploid cells are life’s stress responders as the more than doubling of a cell’s genome has been shown to enable resistance to genotoxic stress. Cells are exposed to various sources of genotoxic stress, including from ultraviolet light. Here we find that ultraviolet light induces DNA damage causing apoptosis and the subsequent generation of giant, multinucleated polyploid cells in the fruit fly epithelium. Unlike in other models studied to date, cell fusion is the predominant response to genotoxic stress and appears to be essential for tissue repair. This study also determines that polyploidization post UV stress is dependent on caspase activation, suggesting a conserved mechanism to initiate cellular multinucleation both in development as well as in response to life’s stressors.

## INTRODUCTION

All organisms undergo various forms of tissue damage throughout their lifetimes from injury, irradiation, and/or toxins. Tissues must compensate for damage-induced cell loss to restore tissue integrity and physiology. In cases where cell division is limited, polyploid cell growth has been shown to be a driver of tissue repair (1, 2). Polyploidy is the more than doubling of a cell’s diploid genome, which can arise by an incomplete cell cycle, known as endoreplication, or by cell fusion. The increase in cell size and DNA content is sufficient to restore tissue mass in several animal tissues studied to date, including the mouse kidney and liver, where cells become polyploid through endoreplication to restore organ mass post-injury (3, 4). Likewise, several tissues in the fruit fly, *Drosophila melanogaster*, rely on polyploidization to compensate for cell loss and restore organ function (5–8). The genetic tractability of the fruit fly has made it a useful model to elucidate the role and regulation of polyploidy in tissue repair.

Polyploid cells arise by endoreplication in most fly tissues studied to date, except the cuticular epithelium, in which polyploid cells are generated by both cell fusion and the endocycle (6, 9–11). Thus, the *Drosophila* epithelium is a beneficial model to discover how stress induces different mechanisms of polyploidization and, in turn, how different ploidy states (i.e., multinucleated versus mononucleated polyploid cells) enable adaptation to a particular stress. The epithelium in the adult fruit fly is composed of a monolayer of post-mitotic diploid cells that, when stressed by a puncture injury, will generate multinucleated, polyploid cells. These polyploid cells are advantageous for tissue repair as they enable healing in the presence of genotoxic stress (12).

Previous studies proved that endocycling cells in the *Drosophila* ovary are resistant to DNA damage caused by gamma irradiation (13). The p53-dependent DNA damage response pathway was epigenetically silenced, hence the polyploid cells could accumulate double-stranded DNA breaks and not undergo apoptosis post-irradiation. Polyploid neurons in the adult fly brain have also been shown to survive UV-induced DNA damage with age (14). Additionally, there is evidence from cancer studies that polyploidy may enable resistance against genotoxic stress (15, 16). However, it is unknown to what extent the different mechanisms of polyploid cell generation (i.e., cell fusion, endocycle, or endomitosis) contribute to genotoxic stress resistance.

Here, we establish an ultraviolet light (UV) injury model in the adult fruit fly where epithelial cells are known to become polyploid in response to stress (6). Irradiating the fly’s ventral abdomen with UV-A (a physiological source of DNA damage) at a low dose induces apoptosis of the post-mitotic diploid cells. While roughly half the cells die, the other cells are able to survive and heal the epithelium within 7 days. Similar to a local injury (i.e., needle puncture wound), the epithelial cells both endoreplicate and fuse, although cell fusion, but not endoreplication, is required for tissue repair following UV-A irradiation. In addition, we find that irradiation-induced apoptosis is required to initiate both cell fusion and the endocycle as expression effector caspase inhibitor, p35, is sufficient to suppress polyploidization. In conclusion, we have found that UV-A irradiation can induce cell fusion generating giant multinucleated, polyploid cells that are essential for cell survival and healing in response to this genotoxic stress.

## RESULTS

### Giant multinucleated, polyploid cells are generated in the fly epithelium in response to UV-A irradiation

To examine how polyploid cells respond and contribute to tissue repair following genotoxic stress in the post-mitotic cells, we used a fruit fly model of UV-A induced injury. To do so, adult fruit flies were irradiated with a low dose (25mJ) of UV-A and the abdominal tissue was dissected, fixed, and immunostained with epithelial-specific antibodies to detect cell junctions (FasIII, a septate junction protein) and epithelial nuclei (GFP, epithelial-specific expression (epi-Gal4>nlsGFP) revealing the epithelial tissue organization in uninjured (0mJ) and days post (dp) UV-A irradiation (Figure 1A). We first validated that UV-A irradiation was sufficient to induce DNA damage by immunostaining for gamma phospho-Histone 2Av (γH2Av), which binds the genome at sites of double stranded DNA breaks (Figure S1) (17). As expected, the uninjured tissue had low, but detectable γH2Av labeled foci in both epithelial and muscle cell nuclei (Figure S1B) (12, 18). By 2dp UV-A, we observed a 2-fold increase in the nuclear γH2Av intensity compared to unirradiated cells (Figure S1B and S1C). The γH2Av expression persisted in the UV-A irradiated epithelial nuclei, suggesting that DNA damage is not repaired even by 10dp UV-A. A high dose (50mJ) of UV-A irradiation was also tested in the adult flies and led to a dosage dependent effect. The tissue showed a higher level of γH2Av within epithelial nuclei at 2dp when compared to the 25mJ treated samples (Figure S1D). However, the 50mJ irradiated flies exhibited more severe tissue damage, which disrupted the tissue during dissection and therefore could not be used for further analysis.

**Figure 1.**
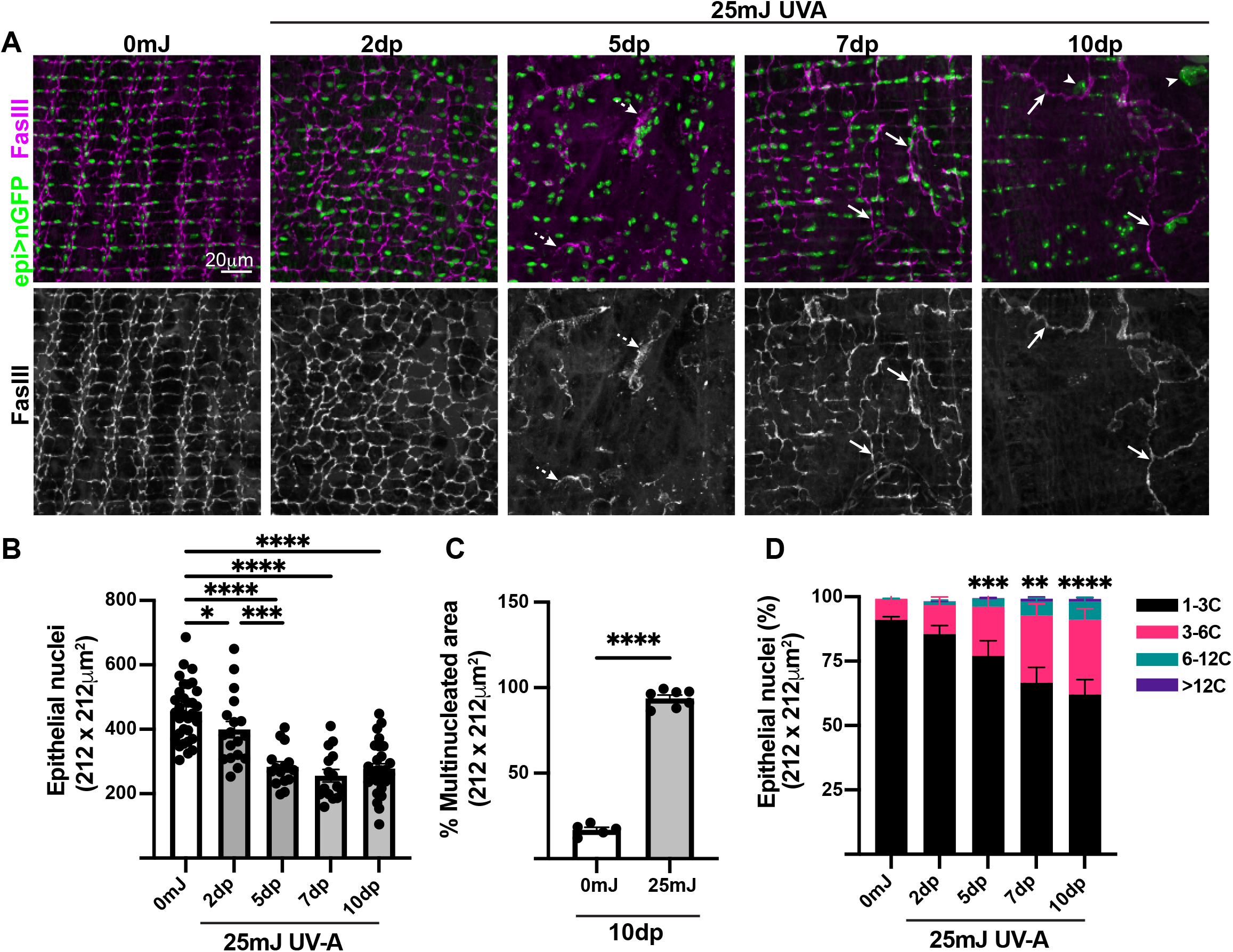
UV-A induced polyploidy in the fly epithelium. (A) Immunofluorescent images from the fly epithelium taken in uninjured (0mJ) and days post (dp) UV-A irradiation (25mJ). Cells are stained for septate junctions (FasIII, magenta) and epithelial nuclei (nGFP, green). Marked are examples of discontinuous (dashed arrows) and continuous (solid arrows) cell junctions and enlarged nuclei (arrowheads). Quantification of epithelial (B) nuclear number, (C) multinucleation at 10dp, and (D) epithelial nuclear ploidy. Data represents the S.E.M. with significance determined by One-way ANOVA with P-value *<0.05, ***<0.001, ****<0.0001.

Next, we examined how UV-A irradiation affected the epithelium in terms of the number of nuclei, multinucleation, and ploidy, as genotoxic stress can lead to apoptosis (Figure 1) (19). The UV-A irradiated epithelium started to show nuclear loss at 2dp and continued to a ∼40% reduction in the number of epithelial nuclei by 5dp. The number of epithelial nuclei then remained steady with no further reduction at 7dp and 10dp, even though nuclear DNA damage (γH2Av nuclear foci) persisted (Figure 1A, 1B, and S1B). The significant reduction in epithelial nuclear number led to a noticeable change in epithelial organization as the septate junctions were disrupted (sparse and discontinuous) by 5dp UV-A (Figure 1A, dashed arrowheads). By 7dp, the epithelial septate junctions were restored and more continuously outlined very large multinucleated cells. The multinucleated cells often extended beyond the image area and so, we quantified the total area occupied by multinucleated cells (see Methods). Only ∼17% of epithelium is multinucleated in *wild-type*, uninjured flies, but by 10dp nearly ∼95% of the epithelium is repaired via generation of multinucleated cells (Figure 1A and 1C).

In addition to the generation of giant, multinucleated cells, we also observed enlarged nuclei by 5-10dp UV-A irradiation. Nuclear size is known to correlate with the DNA content, therefore we measured nuclear ploidy to determine if the epithelial nuclei underwent whole genome duplication (Figure 1A) (20). Epithelial nuclear ploidy was measured using our previously reported method, where nuclear ploidy is determined based on DAPI intensity (21, 22). In doing so, we observed that epithelial nuclei gradually became more polyploid as time progressed post UV-A irradiation (Figure 1D). There was a significant increase in epithelial nuclear ploidy (>3C) by 5dp when ∼40% of the epithelial nuclear number was reduced (Figure 1B and 1D). The epithelial nuclear ploidy further increased in terms of both the percentage of nuclei and total DNA content of each nucleus. At 7-10dp, epithelial nuclear ploidy reached more than 12C with up to 40% of epithelial nuclei exhibiting a >3C DNA content per nucleus. Thus, UV-A irradiation appears to lead to an injury-like response generating both mono- and multinucleated, polyploid cells(6).

### Endoreplication is dispensable for epithelial repair post UV-A irradiation

The increased nuclear ploidy post UV-A indicated that the epithelial cells re-entered an incomplete cell cycle to endoreplicate. To determine when cells entered S-phase, we used the reporter strain, PCNA-EGFP, and conducted a time course post UV-A (Figure S2A) (23). As expected, no PCNA positive epithelial nuclei were observed in the 0mJ control. PCNA+ nuclei gradually peaked at 4dp (∼30, PCNA+ nuclei) and then reduced in number by 6dp (Figure S2B and S2C). Epithelial specific expression was confirmed based on co-localization with the epithelial specific transcription factor, Grh(21, 22). Endoreplication can occur by the endocycle, which bypasses M-phase completely, or by endomitosis in which cells start, but do not finish M phase via failed cytokinesis. To distinguish between the two incomplete cell cycles, the tissues were immunostained with the M-phase marker Cyclin B (CycB). Compared to a positive control, in which *CycB* was overexpressed in the epithelium, there was no detectable CycB expression pre- or post UV-A irradiation (Figure S2D). Therefore, the fly epithelial cells appear to endocycle and not endomitose post UV-A irradiation, leading to whole genome duplication.

Endoreplication contributes to tissue repair as it restores tissue mass and permits healing in the presence of genotoxic stress (12). Thus, we investigated the contribution of endoreplication to epithelial repair post UV-A irradiation, by knocking down *E2F1* in the epithelium using the Gal4/UAS system (*epi>E2F1*^*RNAi*^) (Figure S3). First, we confirmed that *E2F1* knockdown inhibited endoreplication post UV-A irradiation as epithelial nuclear ploidy was significantly reduced and remained in the diploid range (1-3C) for ∼97% of the nuclei (Figure S3A-S3C). Next, we quantified the number of epithelial nuclei pre- and post UV-A irradiation and observed a similar reduction in the nuclear number for both *wild-type* (*epi>+*) and *E2F1*^*RNAi*^ strains (Figure S3D). Similarly, we observed that epithelial cell junctions remained continuous and intact at 7dp UV-A even when endoreplication was inhibited (Figure S3A and S3B). Thus, it appears that endoreplication is dispensable for tissue repair post UV-A irradiation. However, endoreplication may still be necessary to restore tissue mass. Therefore, we measure total tissue ploidy in *wild-type* and *E2F1* knockdown strains pre- and post UV-A irradiation and discovered that endoreplication is necessary to restore loss synthetic capacity, similar to other animal models studied to date (Figure S3E) (5, 6, 21).

### Epithelial cells fuse in response to UV-A irradiation

Next, we examined the role and regulation of the giant, multinucleated epithelial cells that encompassed the epithelium in its entirety post UV-A. Multinucleated cells can arise via endomitosis or cell fusion (1). Since the epithelium is prone to fuse in response to stress and we did not observe expression of the M-phase gene, CycB, we hypothesized that these multinucleated cells were also generated by cell fusion post UV-A stress (6, 24). Cytoplasmic sharing is a reliable means to measure fusion events *in vivo*, so we used the dBrainbow lineage system to mosaically induce expression of the cytoplasmic fluoros EGFP-HSV or EBFP2-HA following Cre-mediated recombination (24, 25). As a result, any bicolored area in the fly epithelium would be indicative of a cell fusion event (Figure 2A). The uninjured (0mJ) epithelium displayed predominantly one color with regions expressing either EGFP or EBFP; while the fluoros in the 7dp UV-A irradiated strains frequently overlapped (Figure 2B). The epithelial area expressing EGFP was ∼45% in the control (0mJ) and significantly increased to ∼85% at 7dp UV-A irradiation (Figure 2C). Comparing the epithelial cell junctions to bicolored areas revealed enlarged cells sharing different levels of EGFP and EBFP expression due to the merging of various number of cells which generated the multinucleated cells (Figure 2B and 2B’-2B’’’). To confirm that the bicolored cells arose by cell fusion and not endoreplication, we generated fly strains where endoreplication could be inhibited (*epi>E2F1*^*RNAi*^). Blocking endoreplication did not affect %EGFP+ area pre- or post UV-A indicating that cytoplasmic sharing of fluoros was primarily due to cell fusion (Figure 2C and 2D).

**Figure 2.**
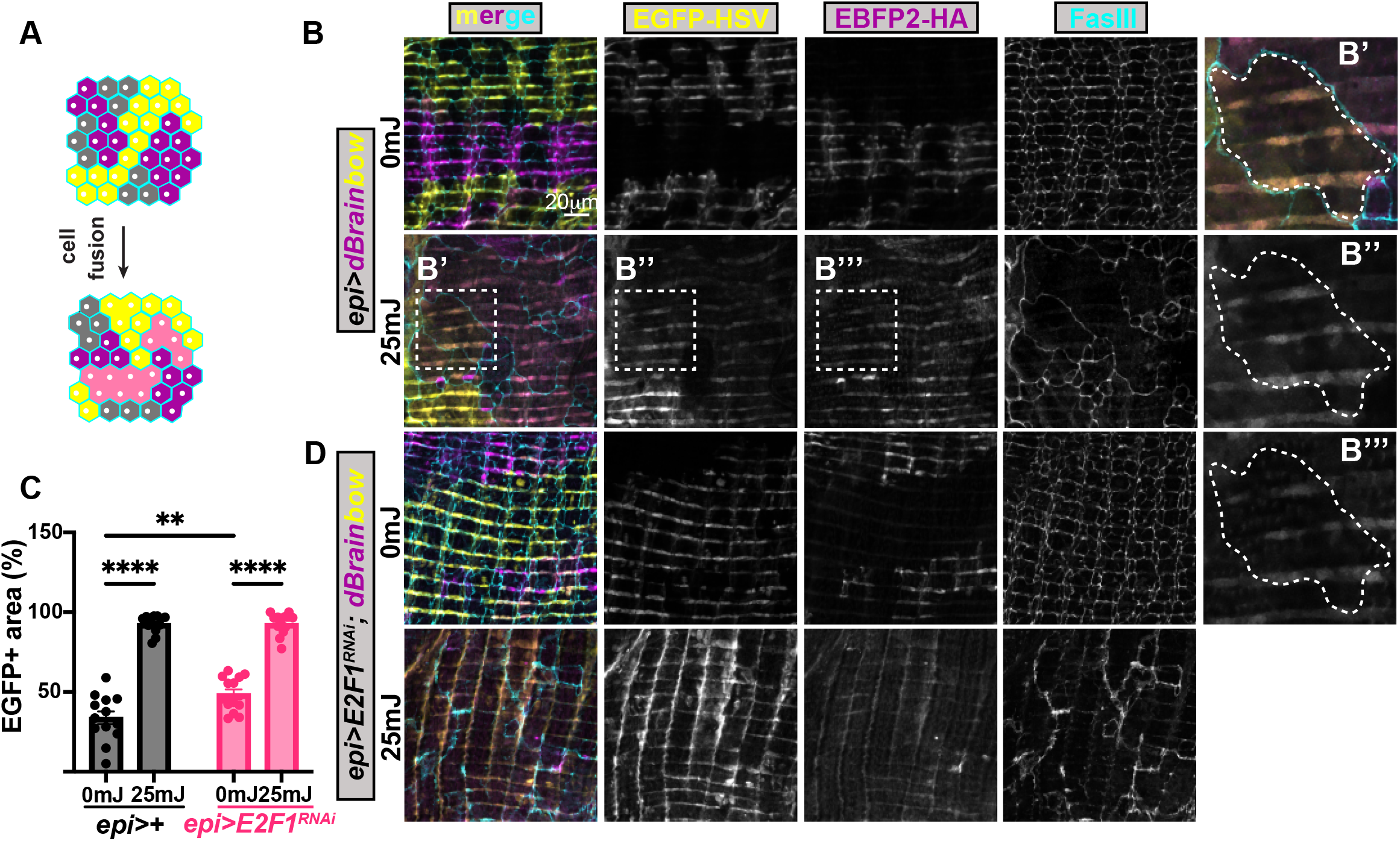
Epithelial cells fuse in response to UV-A irradiation. (A) Illustration of mosaically labeled tissue where cells expressing more than one color are indicative of cell fusion. (B) Immunofluorescent images of the fly epithelium at 7dp +/- UV-A irradiation in *wild-type* dBrainbow strain with fluoros EGFP (yellow) and EBFP2 (magenta). Cell junctions (FasIII, cyan). (B’-B’’’) Zoomed in example of a multinucleated, bicolored cell (dashed white line). (C) Quantification of epithelial area expressing EGFP. Data represents the S.E.M. analyzed by Two-way ANOVA and P-value **<0.01 and ****<0.0001. (D) Immunofluorescent images of the fly epithelium at 7dp +/- UV-A irradiation in dBrainbow, *E2F1*^*RNAi*^ strain.

### Epithelial cell fusion appears to be essential for tissue repair post UV-A

Cell fusion appears to be the prominent response to UV-A irradiation; thus, we tested whether cell fusion is required for tissue repair. Previous studies have shown that epithelial specific expression of dominant negative Rac GTPase (*Rac*^*DN*^) is sufficient to inhibit the generation of multinucleated cells in response to injury or aging (6, 24). Epithelial specific expression of *Rac*^*DN*^ does not affect epithelial development, but after UV-A irradiation only mononucleated epithelial cells were observed on the epithelial periphery (Figure 3A and 3B). The central region of the epithelium was devoid of nuclei in the *Rac*^*DN*^ strain, while big multinucleated cells were observed throughout the epithelium from the *wild-type* strain. Quantifying epithelial nuclear number revealed a significant reduction in total epithelial nuclei, indicating exacerbated apoptosis and a resulting defect in tissue repair (Figure 3C).

**Figure 3.**
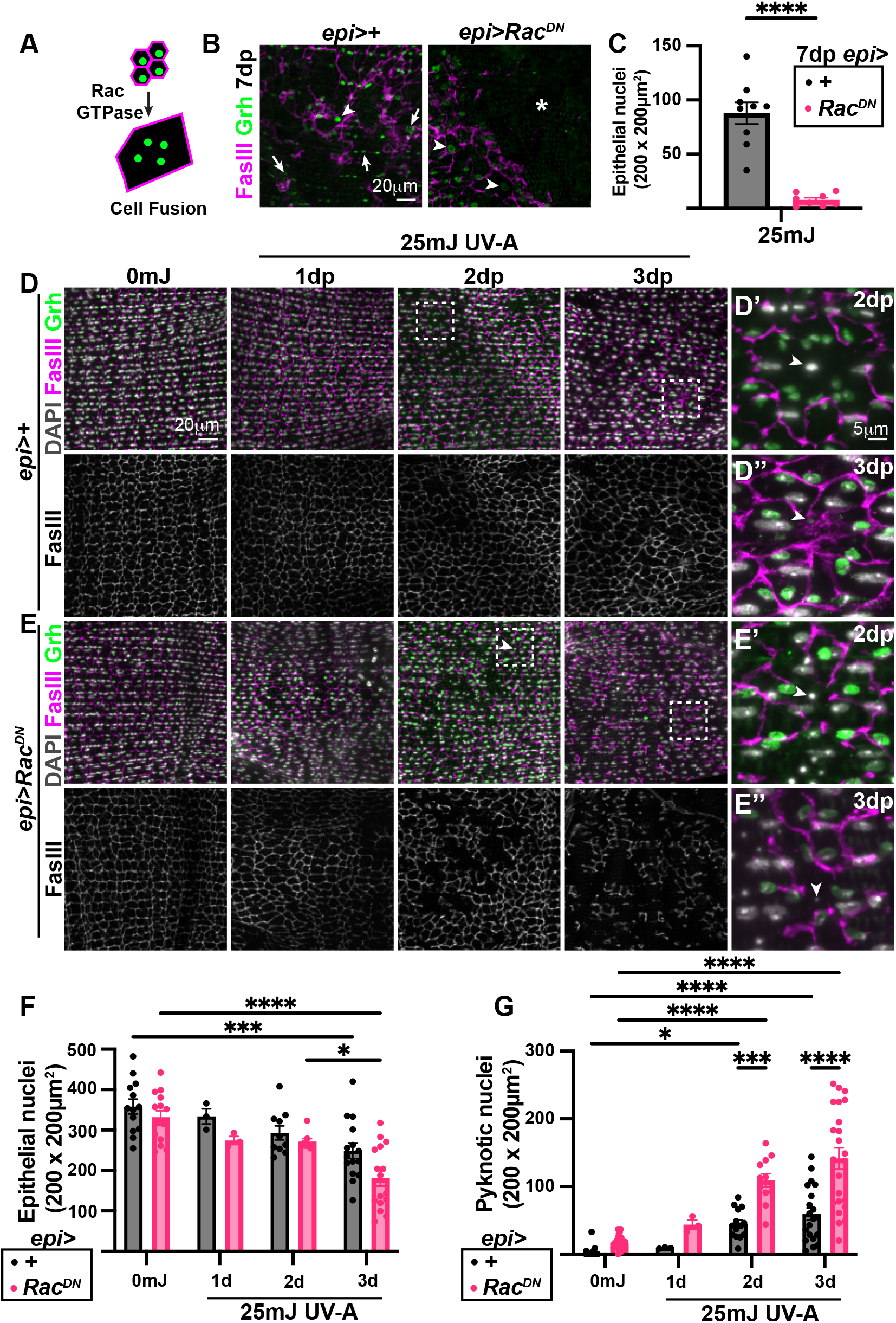
Expression of dominant negative Rac GTPase sensitizes epithelial cells to apoptosis post UV-A irradiation. (A) Model of Rac GTPase dependent cell fusion in the fly epithelial cells. (B) Immunofluorescent images of the fly epithelium at 7dp UV-A irradiation. Epithelial cell junctions (FasIII, magenta) and epithelial nuclei (Grh, green) are shown. Denoted are nuclei within syncytia (arrows) and mononucleated cells (arrowheads), as well as an area devoid of cells/nuclei (*). (C) Quantification of epithelial nuclei number, which is severely reduced post UV-A when dominant negative Rac (*Rac*^*DN*^) is expressed. (D and E) Immunofluorescent images of fly epithelium from uninjured (0mJ) and indicated days post (dp) 25mJ UV-A. Boxed regions show examples of pyknotic nuclei (arrowheads) at 2dp and 3dp UV-A. All nuclei (DAPI, gray). (F and G) Quantification of epithelial nuclear loss and apoptosis, respectively. Data represents the S.E.M. analyzed by Student T-test or Two-way ANOVA and P-value *<0.05, ***<0.001, and ****<0.0001.

Next, we performed a time course at 0, 1, 2, and 3dp UV-A irradiation to determine the cause of the dramatic reduction in the number of epithelial nuclei. To start, we observed no significant difference in epithelial organization at 0 or 1dp UV-A irradiation, as the total number of epithelial nuclei was comparable in both strains and time points (Figure 3D-3F). However, by 2dp UV-A, noticeable changes in the epithelial organization were evident. There were areas devoid of cell junctions and nuclei, suggesting that apoptosis occurred. Zoom-ins (3D’^-^’’ and 3E’^-^’’, boxed regions) revealed fragments, or condensed puncta known as pyknotic nuclei at the center of areas devoid of FasIII septate junction staining. The condensed or fragment nuclei are characteristic of programmed cell death/ apoptosis (26). In most cases, the epithelial nuclei lost their Grh staining but were scored based on their position and nuclear morphology using the DNA intercalating dye, DAPI. The frequency of pyknotic nuclei was further enhanced by *Rac*^*DN*^ expression with several more epithelial areas devoid of cell junctions (Figure 3E). Quantifying the number of pyknotic nuclei revealed that the epithelial cells expressing *Rac*^*DN*^ had substantially more apoptotic nuclei than the *wild-type* cells (Figure 3G). This was further enhanced by 3dp UV-A irradiation as the *Rac*^*DN*^ expressing cells had a greater than 2-fold increase in pyknotic nuclei. As a result, a significant decrease in the number of viable epithelial nuclei was observed by 3dp UV-A in *Rac*^*DN*^ expressing cells (Figure 3F).

The induction of apoptosis and cell fusion appear to occur simultaneously, making it difficult to determine the cause versus the consequence of the epithelial tissue repair defect post UV-A irradiation. The changes in epithelial cell junction morphology suggest that a defect in cell fusion may precede apoptosis as the septate junction protein, FasIII, becomes enriched at the intersection of multiple cell borders in the *wild-type* strain (Figure 3D’ and 3D”). Meanwhile, in the cells expressing *Rac*^*DN*^, there is no indication of FasIII remodeling as the septate junctions appear broken and continue to disintegrate over the time course (Figure 3E’ and 3E”). To further confirm these observations, we thought to identify other regulators of epithelial cell fusion.

Rac GTPase is a member of the Rho family of GTPases, which regulates actin polymerization in cellular development and wound healing (27–29). The Rho family of GTPase have been shown to regulate cell fusion in muscle and fungi, thus we tested if epithelial expression of a dominant negative Cdc42 or Rho would also sensitize epithelial cells to UV-A irradiation (Figure S4) (30–32). The epithelial specific expression of *Cdc42*^*DN*^, similar to *Rac*^*DN*^, resulted in enhanced nuclear loss by 7dp UV-A irradiation (Figure S4). We observed tissue areas devoid of nuclei with mononucleated cells at the borders, while, *Rho*^*DN*^ expressing cells were evenly dispersed across the tissue with the overlaying lateral muscle fibers as visualized by the DAPI stained nuclei (Figure S4A). Expression of *Rho*^*DN*^ did not significantly impact the epithelium as a comparable number of nuclei were lost relative to the *wild-type* (Figure S4B). In conclusion, we have found that Cdc42, like Rac, appears to be required for epithelial cell fusion and tissue repair following UV-A irradiation.

### Caspase-dependent cell death is required for polyploidization post UV-A irradiation

To elucidate the link between cell apoptosis and polyploidy, we blocked cell apoptosis by overexpressing the baculoviral antiapoptotic protein, p35 (*epi>p35*^*OE*^), which counteracts the effector caspases in the apoptotic pathway (33). As compared to the *wild-type* (*epi>+*) strain, *p35*^*OE*^ did not appear to affect the development of the fly epithelium as the cellular organization was normal despite a slight increase in the number of nuclei per unit area (Figure 4A-4C). Following UV-A irradiation, however, *p35*^*OE*^ significantly reduced nuclear loss (Figure 4B and 4C). Fewer enlarged nuclei were also observed in the *p35*^*OE*^ strain, and nuclear ploidy measurements confirmed that the endocycle was inhibited (Figure 4D).

**Figure 4.**
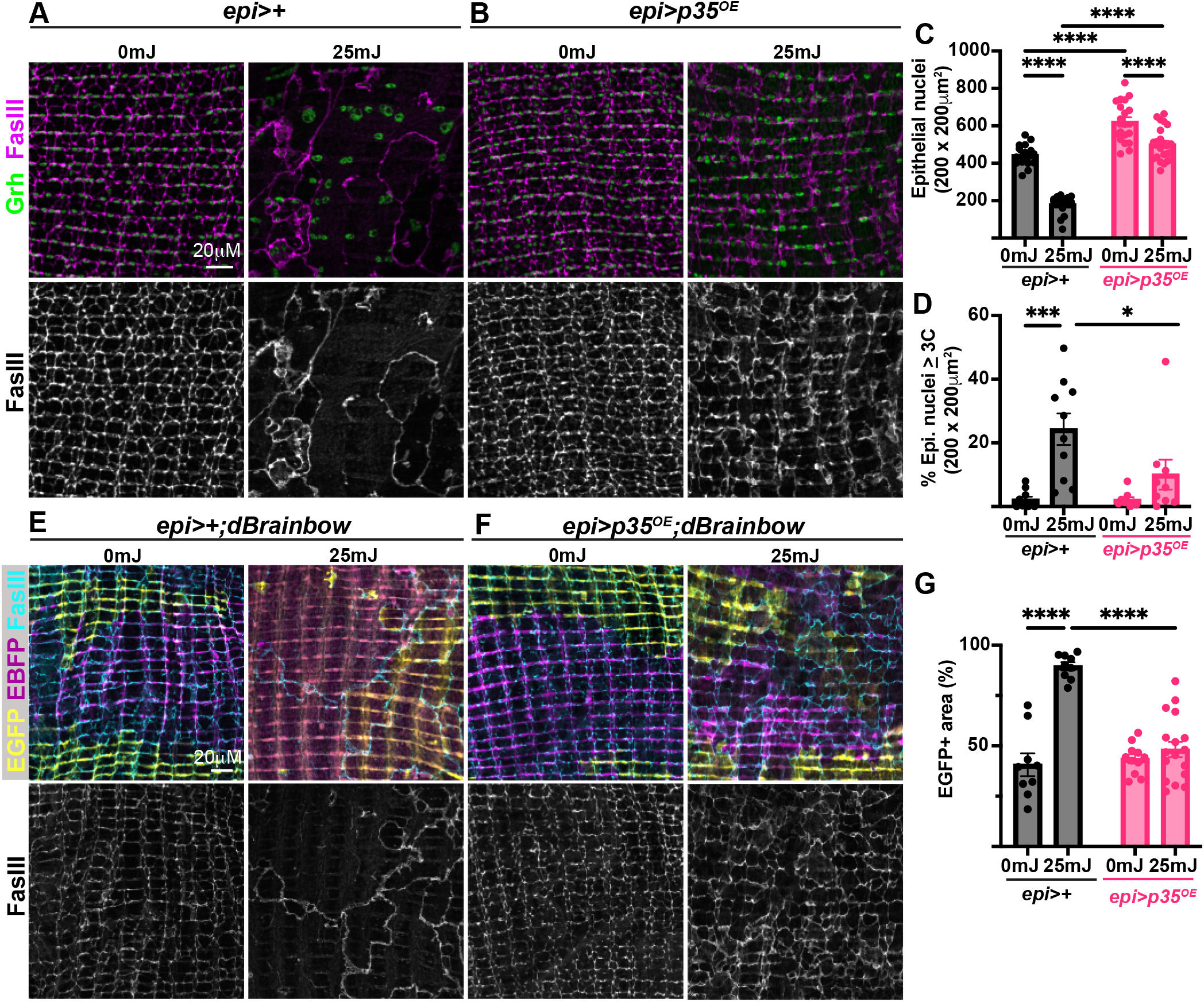
Caspase-dependent apoptosis is necessary for polyploidization post UV-A irradiation. (A and B) Immunofluorescenct images of fly epithelium at 7dp +/- UV-A in the strains indicated. Epithelial nuclei (Grh, green) and cell junctions (FasIII, magenta). Quantification of (C) epithelial nuclei number and (D) ploidy in both strains at 7dp. (E and F) Immunofluorescenct images of fly epithelium at 7dp +/- UV-A in the dBrainbow strains with fluoros EGFP (yellow) and EBFP (magenta). Cell junctions (FasIII, cyan). (G) Quantification of epithelial area expressing EGFP in dBrainbow strains. Data represents the S.E.M analyzed by Two-way ANOVA with P-value *<0.05, ***<0.001, and ****<0.0001.

The epithelial septate junctions were still expressed, but were disordered post UV-A irradiation making it difficult to quantify the extent of multinucleation. Instead, we used dBrainbow mosaic labeling technique to determine whether cell fusion was also inhibited. As shown previously, the *wild-type* strain had extensive overlap between EGFP-HSV and EBFP2-HA fluoros post UV-A (Figure 2B and 4E). The epithelial area expressing EGFP significantly increased to ∼89% at 7dp UV-A compared to 41% in unirradiated (0mJ) *wild-type* (epi>+; dBrainbow) strain (Figure 4G). In contrast, both the uninjured (0mJ) and UV-A irradiated *p35*^*OE*^ strains displayed predominantly one fluoro with regions expressing either EGFP or EBFP (Figure 4F and 4G). Therefore, inhibition of apoptosis through expression of this effector caspase inhibitor is sufficient to strongly reduce cell fusion post UV-A irradiation.

Next, we questioned if it was cell death itself or a caspase-dependent signal that initiated polyploidization. To do so, apoptosis was directly activated by conditionally expressing the pro-apoptotic *Drosophila* gene, *hid* (33–35). We used a Gal80 construct that has the auxin inducible degron to conditionally regulate Gal4 dependent *hid* expression in the fly epithelium for 1d or 3d after the flies eclosed (Figure S5A)(36). This led to a dosage effect, where flies expressing *hid* for 1d lost ∼48% of their epithelial nuclei, whereas flies expressing *hid* for 3d lost ∼87% of their epithelial nuclei (Figure S5B-S5C). Interestingly, giant multinucleated cells were observed in both apoptotic conditions, but the nuclear ploidy was only significantly increased when there was severe nuclear loss in the epithelium (Figure S5C and S5D). As a result, we can conclude that an apoptotic signal is necessary for epithelial cells to fuse, but endoreplication requires a threshold of nuclear loss within the epithelium to be triggered.

## DISCUSSION

Here, we created a fruit fly model of ultraviolet light-induced injury. Exposing the adult fruit flies to a low dose of UV-A was sufficient to cause significant cell loss. The epithelium, in turn, responded by growing instead of dividing the surviving cells. The enlarged epithelial cells predominantly arose by cell fusion and appeared to span the entire tissue area. Unlike with a puncture wound, where cell fusion and endocycling synergize to close the wound, the endocycle appears to be dispensable following UV-A irradiation (6). This is likely because cell fusion provides a more rapid response to widespread damage, accelerating wound closure (1, 6).

### Restoration of tissue mass and activation of the endocycle

Even though endoreplication appears to be dispensable for epithelial wound healing in this UV-A injury model, it is still required to restore tissue mass. Measuring the DNA content (C-value) of the remaining nuclei indicated that the total tissue ploidy was restored with fewer nuclei post UV-A irradiation as the surviving nuclei contained higher ploidy (≥4C). In addition, the endocycle has been shown to enable resistance to genotoxic stress and DNA damage, yet here we find that the endocycle is dispensable (13, 14, 16). Apoptosis, as measured by the number of epithelial nuclei that survive UV-A irradiation, was not changed when the endocycle was inhibited compared to the *wild-type* strain. This may be due to the dual mechanisms of polyploidization in the adult epithelium, as cell fusion, which is the predominant response to UV-A irradiation, appears to be essential to protect the epithelial cells against apoptosis.

DNA damage has been shown to be the initiator of endoreplication for a variety of cell types in response to differentiation, physiological, and pathological cues. This includes mouse mammary gland cells in which endoreplication occurs following replication stress, as well as giant macrophage cells that arise in response to DNA damage from reactive oxygen species (37–39). In these cases, genotoxic stress leads to endoreplication by truncating or bypassing mitosis; however, it remains unknown how DNA damage would result in cell fusion. Here, we suspect it may be through activation of canonical caspase-dependent apoptotic pathway (40).

### The crosstalk between apoptosis and cell fusion

Studies on syncytial cell types, including the skeletal muscle, bone-resorbing osteoclasts, and syncytiotrophoblasts in mammals, have shown that activation of effector caspases, including caspase 3, is necessary for cell-to-cell fusion (41–43). Similarly, we find that overexpression of the effector caspase inhibitor, *p35*, is sufficient to inhibit epithelial cell fusion post UV-A irradiation (44, 45). In osteoclasts, the effector caspase 3 was shown to cleave the La protein, which facilitates cell fusion through phosphatidylserine exposure on the plasma membrane (42, 46). However, it remains to be determined whether this mechanism is conserved in other animal cell types to initiate cell fusion in development or in response to stress.

This fits with our findings, as we were not able to untangle Rac GTPase’s role in apoptosis and fusion post UV-A irradiation. Apoptosis, as observed by pyknotic nuclei, occurred simultaneously with epithelial cell junctional remodeling. Epithelial border breakdown occurred at the site of pyknotic nuclei in both the *wild-type* and *Rac*^*DN*^ expressing epithelial cells. However, the frequency of pyknotic nuclei was significantly enhanced by *Rac*^*DN*^ expression, suggesting that Rac inhibits apoptotic signaling in the process of regulating cell fusion in this model.

Interestingly, overexpression of the effector caspase inhibitor, p35, was also sufficient to inhibit endoreplication post UV-A irradiation. The adult *Drosophila* epithelial cells are post-mitotic and do not express M-phase genes (6, 12). Therefore, the epithelial cells are unlikely to experience replication stress that would truncate the mitotic cell cycle (16). Instead, previous studies have shown that the degree of nuclear ploidy is dependent on the tissue’s cellular density, which is regulated by the conserved Hippo-Yki signal transduction pathway (21). As a result, we suspect that it is apoptosis itself that is necessary for endoreplication, as severe nuclear loss was needed to increase nuclear ploidy when cell death was genetically activated through *hid* expression. And in the case of cell fusion, it is more likely to be dependent on a non-apoptotic role of caspases as observed in cells, such as osteoclasts, that undergo cell fusion during development.

### Cell fusion as a stress response

While cell fusion is known to be required for the development of a number of multinucleated cells, its role and regulation as an adaptation to stress remains more poorly understood (47). Multinucleated cells, also known as syncytia, have been reported to arise most commonly upon viral infection due to an encoded fusogen (48). There are also reports that cell fusion contributes to the generation of giant polyploid cancer cells (49, 50). However, most mammalian tissues are complex, and a common challenge is to distinguish whether the multinucleated cells identified arose by failed cytokinesis or cell fusion.

The fruit fly, *Drosophila*, has emerged as a model to study how stress triggers polyploidization, particularly in epithelial cell fusion. The epithelial cells underlying the cuticle exoskeleton have been found to undergo cell fusion at all post-embryonic stages of life (i.e. larval, pupa, and adult) post-injury and with age (6, 9, 10, 24, 51). Here, we have identified another stress that induces cell fusion, UV-A irradiation. Utilizing *Drosophila*’s extensive genetic toolkit, the extent of cell fusion was determined through expression of the dBrainbow mosaic labeling system. A week post UV-A irradiation led to near complete overlap of the dBrainbow fluoros, whereas before irradiation, the fluoros rarely overlapped. Inhibiting effector caspase activity with p35 kept fluoros separate, similar to the unirradiated flies. The dBrainbow technique was essential to validate whether cell fusion was inhibited by p35 expression, as the septate junctions became disorganized post UV-A irradiation complicating the analysis.

Lastly, we determined that cell fusion post UV-A irradiation is dependent on the Rho family GTPases, Rac and Cdc42, which play a conserved role in cell fusion. Both Rho GTPases regulate actin polymerization, an essential process in cell fusion from fungal species to the development of syncytial muscle cells (30–32). Epithelial cells on the periphery of the UV-A induced injury remained mononucleated when either Rac or Cdc42 dominant negative mutants were expressed. Thus, their GTPase activity appears to be necessary for cell fusion post UV-A irradiation, although we cannot rule out additional roles for Rac or Cdc42 in epithelial cell survival or wound healing in this model. While the molecular mechanism of epithelial cell fusion in response to stress remains to be identified, here we discover a link between apoptosis and cell fusion, demonstrating that an apoptosis-dependent mechanism is necessary for polyploidization post UV-A irradiation.

## METHODS AND MATERIALS

### Fly husbandry and strains

All *Drosophila melanogaster* strains and crosses used in these experiments were kept in vials containing a standard corn syrup and soy food (Archon Scientific) at 25°C, 60% humidity, and a 12-hour light/dark cycle. See Table S1 for a complete list of strains used in this study. All crosses were created using standard mating schemes in *Drosophila*, beginning with 10 virgin females and 5 males. The Gal4/UAS system was used to express the dominant negative genes and gene-specific RNAi using an epithelial-specific GMR51F10-Gal4 driver (6, 21).

### UV-A irradiation

Flies were collected, aged to 3-days old, and anesthetized using Fly-Nap (Caroline Biologicals, cat#173010) for 3 minutes to ensure they remained asleep during UV-A irradiation. Once asleep, the *Drosophila* strains were immobilized with their ventral abdomen facing up using double-sided tape and irradiated at 4cm from lamps in the Fisherbrand Stratalinker with UV-A bulbs using a dose of 25mJ. The flies were returned to food vials and left to recover for the indicated times in days post (dp) irradiation.

### *Drosophila* dissection and immunostaining

*Drosophila* abdominal dissection and immunostaining were performed on female flies as previously reported (22). The fly abdomens were immobilized on a Sylgard plate, then fixed in 4% paraformaldehyde for 30 minutes, washed, and permeabilized with 1x PBS, 0.3% Triton X-100, and 0.3% BSA. The samples were then stained overnight in primary followed by secondary antibody solutions. See Table S2 for a complete list of all antibodies used in this study. Lastly, all samples were stained with DAPI (1:5000) to visualize all nuclei.

### Fly imaging and analysis

The fixed, stained, and mounted abdomen samples were imaged using a Zeiss AxioImagerM2 microscope with an Apotome and a 40X dry objective with a Hamamatsu Orca-Flash 4.0 camera. Samples for Figures 1, 2, and 4 were imaged using the 40× oil (Zeiss Immersol 514F) objective on a Zeiss LSM 880 Airyscan. Z-stack images were taken with a slice size of 0.5μm. Using ImageJ/FIJI, images were flattened into MAX intensity stacks for all except the DAPI stain, which was flattened as a SUM intensity stack, and then merged to generate the composite images that are shown.

### Nuclei Counting

Images were rotated and cropped to 100 x 100μm^2^, 200 x 200μm^2^, or 212 x 212µm^2,^ as indicated on the y-axis of the figure graphs. The cell counter tool in ImageJ/FIJI was used to quantify the viable nuclei, which colocalized with epithelial-specific Grh or nuclear GFP and DAPI stains. Nuclei that overlapped with the image border were excluded from the analysis. Apoptosis leads to condensation and fragmentation of nuclei, generating pyknotic nuclei. Pyknotic nuclei were identified as bright DAPI fragments or spots in the epithelial row that did not co-stain with Grh. The number of pyknotic nuclei may be higher than viable nuclei loss from tissue damage, as nuclei often fragment into multiple pieces during cell death.

### Ploidy Analysis

Epithelial nuclear ploidy was measured as previously reported (22). Using FIJI, regions of interest (ROIs) were outlined based on epithelial specific nuclear GFP or Grh and mapped to the SUM projection of the Z-stack DAPI image to measure the DAPI intensity. Overlapping nuclei were excluded from the analysis. The raw DAPI intensity for each nucleus was measured by subtracting the background signals. The uninjured epithelial nuclei served as the diploid (2C) control to calculate the nuclear ploidy number based on the average from at least three samples per condition. Nuclei were grouped based on ploidy class of 1-3C, 3-6C, 6-12C, and >12C.

### Statistical Analysis

Each experiment was repeated in at least duplicate with the indicated replicate number (n) for each *Drosophila* strain in the Source data file. Statistical analysis was performed using Microsoft Excel and GraphPad Prism software. The tests conducted included Tukey’s One-way/Two-way ANOVA and Student T-test with P-values as follows: * (p<0.05), ** (p<0.01), *** (p<0.001), and **** (p<0.00001).

## Data Availability Statement

The authors affirm that all data necessary for verifying the conclusions are present within the article, figures, and Supplemental Information. All raw data points reported in figures can be found in the Source data file. Additionally, the *Drosophila* strains and antibodies used in this study are listed Table S1 and S2 and are available upon request or as referenced.

## Acknowledgments

We would like to thank the members of the Losick lab, including Lydia Bischoff, Stefania Bonnani, Loiselle Gonzalez-Baez, Sophie Jalkut, and Jack Weidenbach for critical review of this manuscript. The fly strains used in this study were purchased from (with support by): the Bloomington Drosophila Stock Center (NIH P40OD018537) and Vienna Drosophila Resource Center. The FasIII antibody used in this study was purchased from the Developmental Studies Hybridoma Bank (created by the NICHD and maintained at The University of Iowa, Department of Biology, Iowa City, IA 52242, USA). Images were acquired using microscope supported by the National Science Foundation under Grant No. 1626072 with additional thanks to Bret Judson whom directs the Boston College Imaging Core for infrastructure and support.

## Funding

This work was supported by Boston College and the National Institute of General Medical Sciences of the National Institutes of Health under Award Number R35GM124691 to V.P.L. The content is solely the responsibility of the authors and does not necessarily represent the official views of the NIH.

## Notes

### Competing Interest Statement

The authors have declared no competing interest.

