## Supplemental Information and Figures for "Epithelial cell fusion is required for tissue repair following UV-A irradiation"

Vicki P. Losick

#### **This PDF file includes:**

Figures S1 to S5

Tables S1 and S2

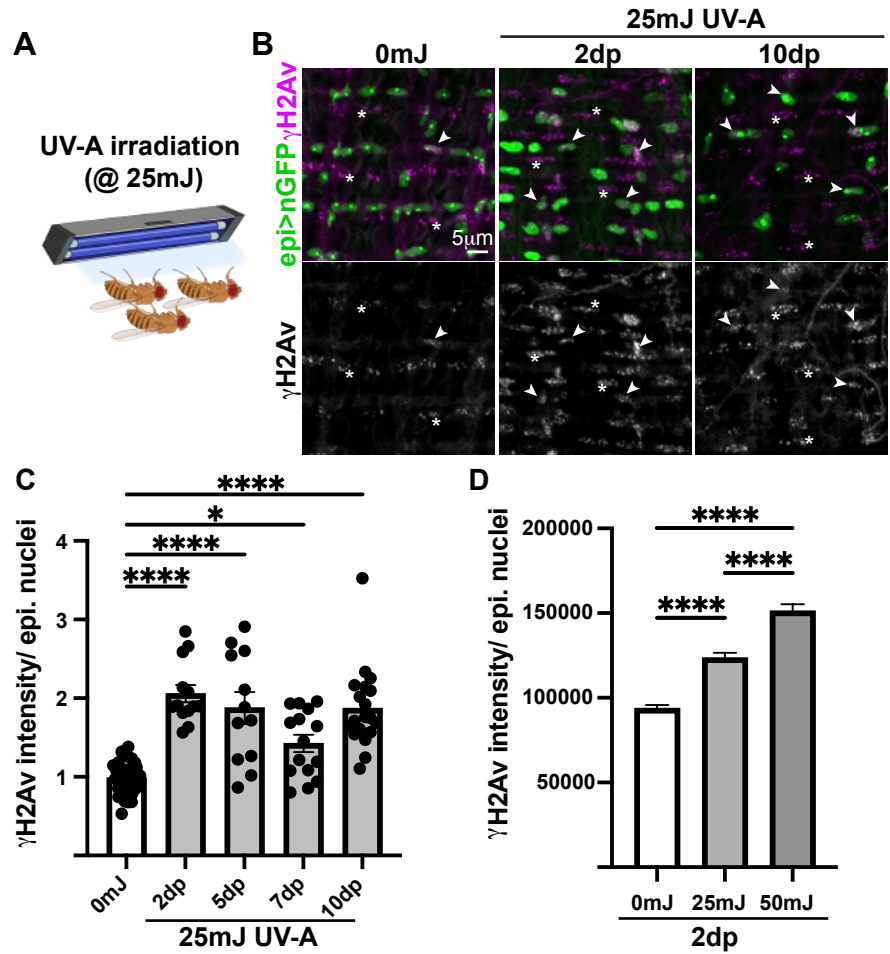

**Figure S1. UV-A irradiation leads to persistent DNA damage in the epithelial cells.**

(A) Adult female flies were irradiated with 25mJ (+) of UV-A to determine how the epithelium responded. (B) Immunofluorescent images of epithelium +/- UV-A at 2 and 10 days post (dp). Epithelial nuclei (Grh, green) and DNA damage (γH2Av, magenta). Examples of epithelial nuclei with DNA damage (arrowheads) and muscle nuclei (\*) in fly abdomen stain with γH2Av regardless of irradiation. (C) Quantification of extent of DNA damage based on γH2Av intensity per epithelial nuclei at time indicated. (D) Quantification of dose-dependent DNA damage based on γH2Av signal per epithelial nuclei at 2dp. Data represents the S.E.M analyzed by One-way ANOVA with P-value \*$<0.05$, \*\*\*\*$<0.0001$.

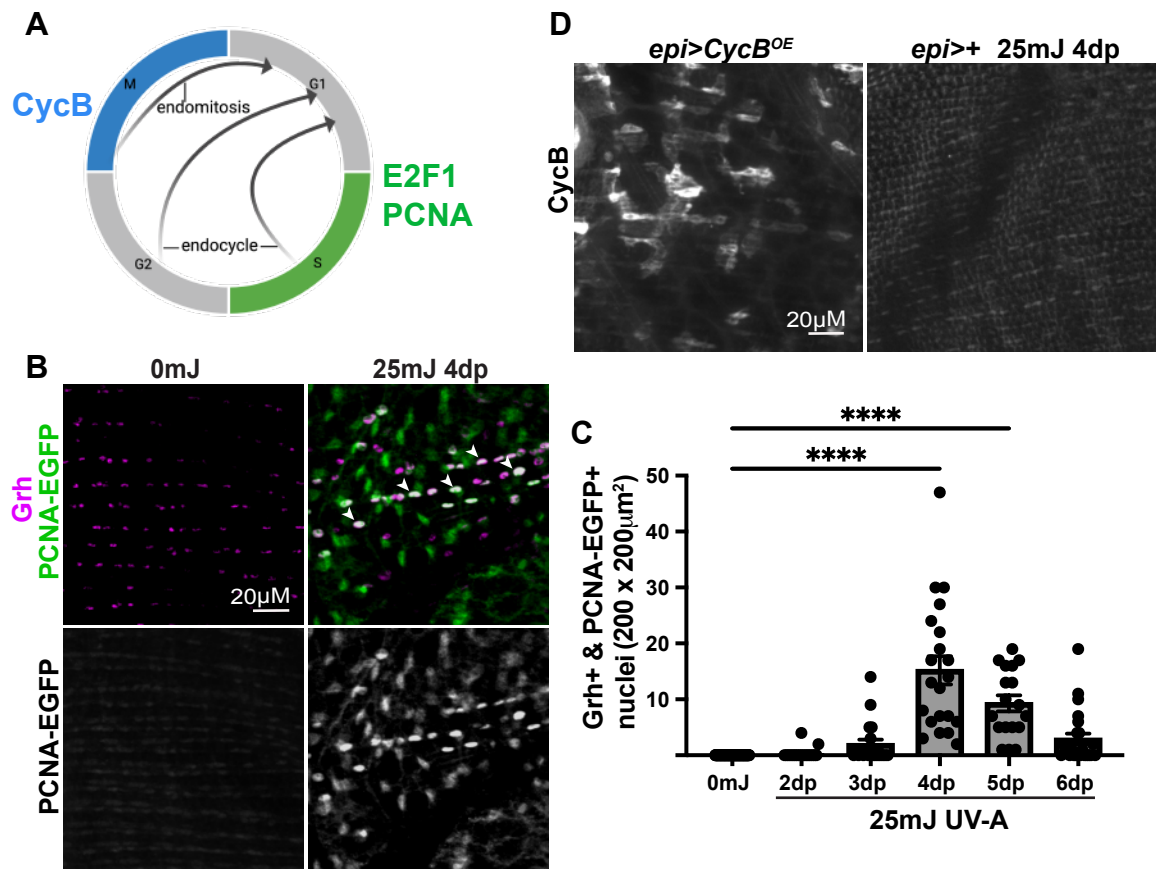

**Figure S2. Fly epithelial cells endocycle at 4d post UV-A.** (A) Illustration of endoreplication, where endocycle bypasses M-phase, but endomitosis truncates M-phase. PCNA is expressed in both incomplete cell cycles, but CycB is specific to endomitosis. (B) Immunofluorescent images of PCNA-EGFP expression at 0dp and 4dp UV-A irradiation. Epithelial nuclei (Grh, magenta) and S-phase marker (PCNA-EGFP, green). (C) Quantification of epithelial/PCNA+ nuclei indicates that the fly epithelial cells enter the endocycle at 4dp UV-A. Data represents the S.E.M analyzed by One-way ANOVA with P-value \*\*\*\*<0.0001. (D) Immunofluorescent images of fly epithelium where CycB is only detectable when overexpressed (*epi>CycB<sup>OE</sup>*) as positive control. CycB was not detected post UV-A irradiation at 4dp.

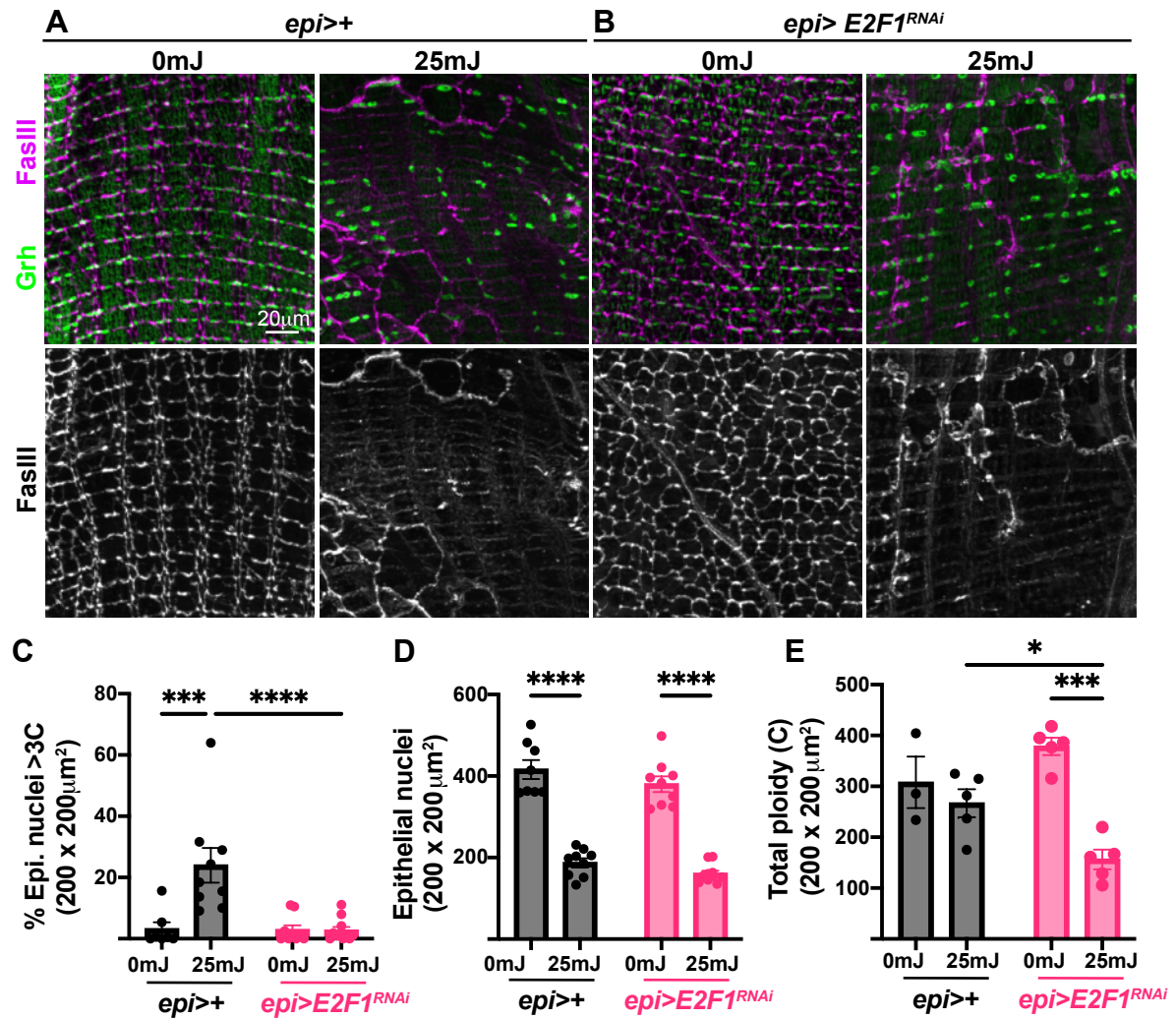

**Figure S3. Epithelial endocycling is dispensable for tissue repair, but essential to restore total tissue ploidy post UV-A.** (A and B) Immunofluorescent images of fly epithelium at 7dp +/- UV-A . Epithelial cell junctions (FasIII, magenta) and epithelial nuclei (Grh, green) are shown. (C) Quantification of epithelial nuclear ploidy, (D) nuclear number, and (E) total tissue ploidy in *wild-type* and *E2F1<sup>RNAi</sup>* strains. Data represents the S.E.M analyzed by Two-way ANOVA with P-value \*<0.05, \*\*\*<0.001, and \*\*\*\*<0.0001.

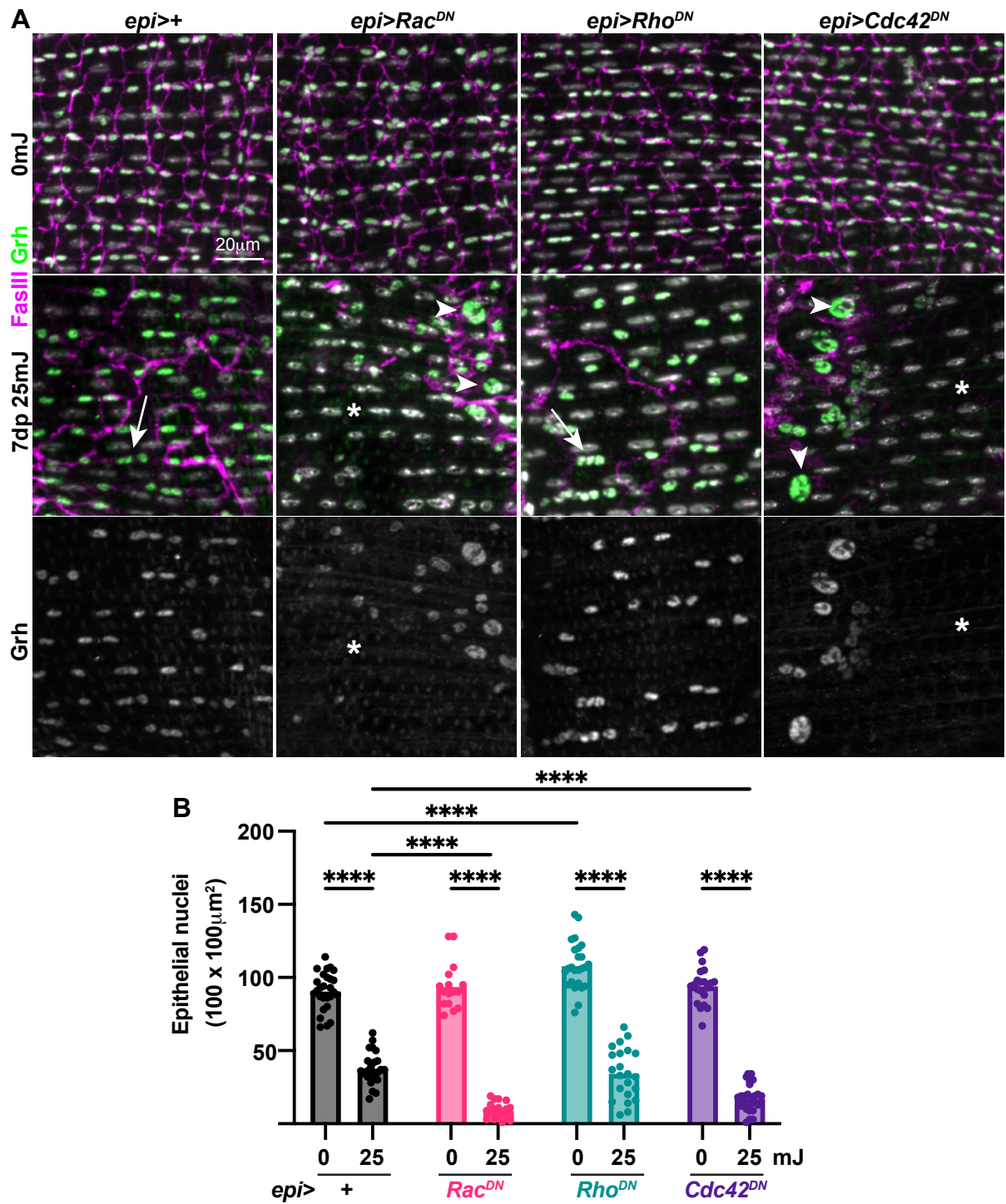

**Figure S4. Both Rac and Cdc42 are required for cell survival and fusion post UV-A irradiation.**

(A) Immunofluorescent images of the fly epithelium +/- UV-A in strains indicated. Epithelial cell junctions (FasIII, magenta), epithelial nuclei (Grh, green) and all nuclei (DAPI, gray). Denoted are nuclei within syncytia (arrows), mononucleated cells (arrowheads), and areas devoid of all cells (\*). (B) Quantification of epithelial nuclei number, which is severely reduced post UV-A when dominant negative *Rac* or *Cdc42* (*Rac<sup>DN</sup>* and *Cdc42<sup>DN</sup>*) are expressed. Data represents the S.E.M. analyzed by Two-way ANOVA and P-value \*\*\*\*<0.0001.

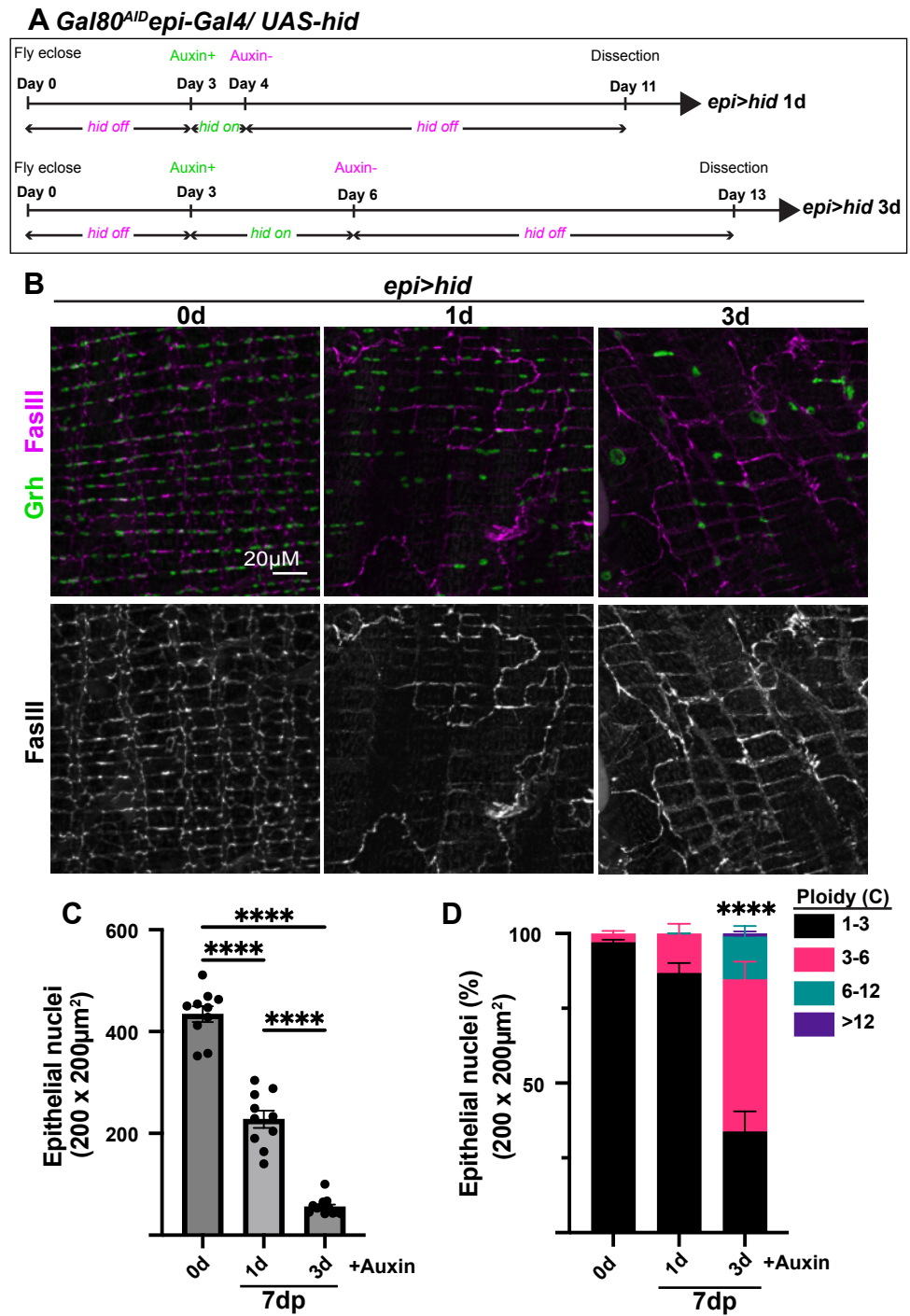

**Figure S5. Genetic induction of apoptosis is sufficient to induce polyploidization in the fly epithelium.** (A) Illustration of the conditional regulation of the apoptotic gene, *hid*, in the fly epithelium using Gal4 inhibitor Gal80 under control of the auxin-inducible degron. Without auxin in fly food, Gal4 is inhibited by Gal80 and *hid* expression is off, whereas addition of auxin to the fly food results in degradation of Gal80 and expression of the apoptotic gene for the duration in which flies are fed auxin. (B) Immunofluorescent images of the fly epithelium at 7dp without (0d) or with auxin for times indicated. Epithelial cell junctions (FasIII, magenta) and epithelial nuclei (Grh, green). (C) Quantification of epithelial nuclei number and (D) nuclear ploidy shows dosage-dependent effect due to ectopic expression of *hid*. Data represents the S.E.M. with significance determined by One-way ANOVA with P-value \*\*\*\*<0.0001.

**Table S1. *Drosophila* strains used in the study.**

| Strain | Strain (Genotype) | Stock number | Source |
| --- | --- | --- | --- |
| *w*<sup>1118</sup> | w[1118] | 5905 | Bloomington Drosophila Stock Center |
| *EGFP-PCNA* | w;; EGFP-PCNA<br>attP2/TM3, sb, {hs-hid} |  | *Blythe and Wieschaus.*<br>(2016) |
| *UAS-Rac*<sup>DN</sup> | y[1] w[*]; P{w[+mC]=UAS-<br>Rac1.N17}1 | 6292 | Bloomington Drosophila Stock Center |
| *UAS-dBrainbow* | w[1118]; P{y[+t7.7]<br>w[+mC]=UAS-<br>Brainbow}attP2 | 34513 | Bloomington Drosophila Stock Center |
| *UAS-p35*<sup>OE</sup> | w[*]; P{w[+mC]=UAS-<br>p35.H}BH2 | 5072 | Bloomington Drosophila Stock Center |
| *UAS-Flp* | w[*]; P{w[+mC]=UAS-<br>FLP1.D}JD2/TM3, Sb[1] | 4540 | Bloomington Drosophila Stock Center |
| *epi-Gal4* | w[1118]; P{y[+t7.7]<br>w[+mC]=GMR51F10-<br>GAL4}attP2 | 38793 | Bloomington Drosophila Stock Center |
| *9F6FO* | P{Ubi-<br>p63E(FRT.STOP)Stinger}<br>9F6/CyO | 32250 | Bloomington Drosophila Stock Center |
| *UAS-E2F1*<sup>RNAi</sup> | P{KK100304}VIE-260B | 108837 | Vienna Drosophila Resource Center |
| *UAS-Cdc42*<sup>DN</sup> | w[*]; P{w[+mC]=UAS-<br>Cdc42.N17}3 | 6288 | Bloomington Drosophila Stock Center |
| *UAS-Rho1*<sup>DN</sup> | w[*]; P{w[+mC]=UAS-<br>Rho1.N19}2.1 | 7328 | Bloomington Drosophila Stock Center |
| *UAS-hid* | w[*];P{w[+mC]=UAS-hid.Z}2/CyO | 65403 | Bloomington Drosophila Stock Center |
| *Gal80<sup>AID</sup>* | w[1118]; PBac{y[+mDint2]w[+mC]=tubP-TIR1-2A-GAL80.AID}VK00040 | 92470 | Bloomington Drosophila Stock Center |

**Table S2. Antibodies used in the study.**

| **Antibody** | **Dilution** | **Catalog number**<br>**(Antibody Registry)** | **Source** |
| --- | --- | --- | --- |
| Rabbit anti-Grh | 1:300 | (AB_2568305) | *Losick et al.* 2016 |
| Chicken anti-GFP | 1:200 | A10262(AB_2534023) | Thermofisher |
| Mouse anti-FasIII | 1:50 | 7G10 (AB_528238) | DSHB |
| Rabbit anti-gH2Av | 1:1000 | 600-401-914<br>(AB_828383) | Rockland |
| rabbit anti-GFP | 1:2000 | A11122 (AB_221569) | Thermofisher |
| Rat anti-HA | 1:100 | 11867423001(AB_390918) | Roche |
| Donkey anti-rabbit 488 | 1:1000 | A21206 (AB_2535792) | Thermofisher |
| Donkey anti-rabbit 568 | 1:1000 | A10042 (AB_2534017) | Thermofisher |
| Goat anti-mouse 488 | 1:1000 | A10680 (AB_2534062) | Thermofisher |
| Goat anti-mouse 568 | 1:1000 | A11031 (AB_144696) | Thermofisher |
| Goat anti-mouse 633 | 1:1000 | A21146 (AB_2535782) | Thermofisher |
| Goat anti-rat 568 | 1:1000 | A-11077 (AB_2534121) | Thermofisher |
